# A synthetic genetic array screen for interactions with the RNA helicase *DED1* during cell stress in budding yeast

**DOI:** 10.1101/2022.08.15.504003

**Authors:** Sara B. Carey, Hannah M. List, Ashwin Siby, Paolo Guerra, Timothy A. Bolger

## Abstract

During cellular stress it is essential for cells to alter their gene expression to adapt and survive. Gene expression is regulated at multiple levels, but translation regulation is both a method for rapid changes to the proteome and, as one of the most energy-intensive cellular processes, a way to efficiently re-direct cellular resources during stress conditions. Despite this ideal positioning, many of the specifics of how translation is regulated, positively or negatively, during various types of cellular stress remain poorly understood. To further assess this regulation, we examined the essential translation factor Ded1, an RNA helicase that has been previously shown to play important roles in the translational response to cellular stress. In particular, *ded1* mutants display an increased resistance to growth inhibition and translation repression induced by the TOR pathway inhibitor, rapamycin, suggesting that normal stress responses are partially defective in these mutants. To gain further insight into Ded1 translational regulation during stress, synthetic genetic array analysis was conducted in the presence of rapamycin with a *ded1* mutant and a library of non-essential genes in *S. cerevisiae* to identify positive and negative genetic interactions in an unbiased manner. Here we report the results of this screen and subsequent network mapping and GO-term analysis. Hundreds of candidate interactions were identified, which fell into expected categories, such as ribosomal proteins and amino acid biosynthesis, as well as unexpected ones, including membrane trafficking, sporulation, and protein glycosylation. Therefore, these results provide several specific directions for further comprehensive studies.

## Introduction

During adverse extracellular conditions, such as nutrient deprivation or oxidative stress, cells must re-orient their gene expression profiles to slow growth, conserve resources, and respond to the stressor (Pakos-Zebrucka et al. 2016; Saxton and Sabatini 2017). Protein translation is both highly energy-intensive and a direct determinant of the cellular proteome; thus, it is a natural point of regulation in stress responses (Crawford and Pavitt 2019; Liu and Qian 2014). Indeed, translation undergoes massive reprogramming during stress, wherein bulk translation is repressed, but translation of select “stress-response” mRNAs is upregulated (Gerashchenko et al. 2012; Ingolia et al. 2009). However, the mechanisms underlying this specificity remain incompletely understood.

In budding yeast, *DED1* encodes an essential RNA helicase of the DEAD-box protein family, which are critical for modulating RNA-RNA and RNA-protein interactions throughout gene expression (Valentini and Linder 2021). Its human ortholog, DDX3X, has been implicated in multiple cancers, including frequent mutations in medulloblastoma, and DDX3X mutations also cause an autism-like cognitive disorder (Mo et al. 2021; Northcott et al. 2012; Snijders Blok et al. 2015; Tang et al. 2021). The primary function of Ded1 is thought to be in translation initiation. In normal, pro-growth conditions, Ded1 promotes initiation by unwinding secondary structure in the 5’ UTR of mRNAs and stimulating pre-initiation complex assembly (Gupta et al. 2018; Sen et al. 2015; Sharma and Jankowsky 2014). Furthermore, *ded1* mutation preferentially affects mRNAs with structured 5’ UTRs and increases the utilization of alternative translation initiation sites in target mRNAs (Guenther et al. 2018; Sen et al. 2015).

Interestingly, Ded1 also plays roles in repressing translation. *DED1* overexpression inhibits translation and cell growth, and Ded1 affects the formation of stress granules, stress-dependent, cytoplasmic accumulations of RNA and proteins (Aryanpur et al. 2022; Beckham et al. 2008; Hilliker et al. 2011). Notably, we recently showed that Ded1 mediates the translational response to TOR inactivation (Aryanpur et al. 2019). Specifically, a *ded1* mutant lacking its C-terminal region (*ded1-ΔCT*) was resistant to growth inhibition and translation repression caused by the TOR inhibitor rapamycin. The C-terminal region of Ded1 interacts with the scaffolding factor eIF4G1, and further analysis suggested that eIF4G1 mediates the effects of Ded1 in these conditions (Aryanpur et al. 2019; Hilliker et al. 2011). We proposed a model wherein Ded1 represses translation during TOR inactivation by promoting eIF4G1 dissociation from translation complexes and its subsequent degradation. How this mechanism is regulated and which downstream processes are affected remains unknown, however.

To begin to address these questions, we conducted a synthetic genetic array (SGA) screen with the *ded1-ΔCT* mutant, taking advantage of its resistance to rapamycin-mediated growth inhibition to identify both positive and negative synthetic interactions from the yeast deletion library of non-essential genes (Tong et al. 2001). This screen identified a large number of synthetic interactions with associated GO-terms that include translation, vesicle trafficking, amino acid metabolism, and signal transduction. These hits likely represent upstream regulators, direct interactions, and downstream targets of Ded1 as well as related processes. Compelling candidates will be examined in future studies.

## Methods & Materials

### Screen Design

The yeast strain used in this SGA screen is TBY174 (*MAT*α *his3*Δ*1 leu2*Δ*0 ura3*Δ*0 can1*Δ*0∷P*_*GAL1*_*-T*_*ADH1*_*-P*_*MFA1*_*-spHIS5 lyp1*Δ*0 ded1-ΔCT::Hygro*), which was constructed from the strain Y15583-13.2b from a previous study using similar techniques (Singh et al. 2009). The C-terminal portion of DED1 was removed from Y15583-13.2b and replaced with a hygromycin resistance cassette using the plasmid pUG75 as a template (protocol adapted from (Hegemann and Heick 2011)). The resulting strain was verified using PCR amplification, growth assays, and western blotting. This “query strain” (TBY174) was then used in a standard synthetic genetic array protocol (Singh et al. 2009; Tong and Boone 2006). The yeast knockout library is commercially available and contains approximately 5,000 non-essential gene knockout strains (Horizon Discovery). These strains are constructed using a G418 resistance cassette and have been independently validated and certified by the supplier. The mating type of the strains used are MATa allowing them to be mated to the query strain directly.

The screen procedure can be summarized as: re-array of the knockout library, mating of query strain to the library, selection of zygotes, sporulation, selection of haploids, and growth analysis. To re-array the library, the 96-well plates of the library stock were pinned using a RoToR HDA Pinning robot (Singer) to a 384-well format compatible with the robot. These plates contained solid YPD (10 g/L yeast extract, 20 g/L peptone, 20 g/L agar, 200 µl/L 10N NaOH, 2% glucose) + G418 (200 mg/L) media to maintain selection, and yeast were grown for 48 hours at 30°C. To prepare for mating, the query strain was simultaneously grown in liquid culture for 24 hours, plated to a 384-well format, and allowed to grow for an additional 24 hours at 30°C. The query strain was then replica plated to 14 fresh plates containing YPD. Next, the re-arrayed knockout library plates were replica plated directly on top of the query strain and allowed to grow for 24 hours at 30°C. To select for zygotes, the mated strains were replica plated to new plates containing YPD + G418 + hygromycin (200 mg/L) and allowed to grow for 48 hours at 30°C. For efficient sporulation, the zygote plates were replica plated to plates containing sporulation media (20 g/L agar, 10 g/L potassium acetate, 1 g/L yeast extract, 0.5 g/L glucose, and 0.1 g/L amino acid sporulation supplement; supplement consisted of 2 g histidine, 10 g leucine, 2 g lysine, and 2 g uracil) and allowed to grow for 5 days at 22°C.

Following the sporulation period, the desired double knockout (DKO) haploid strains were isolated using a progression of three different selections. First, MATa sporulated strains were selected by replica plating onto Singer-compatible plates containing SD media (20 g/L agar, 20 g/L glucose, 1.7 g/L yeast nitrogen base without ammonium sulfate and amino acids, 1 g/L monosodium glutamic acid, and 2 g/L amino acid supplement; supplement consisted of 1.2 g adenine, 1.8 g isoleucine, 3.6 g leucine, 1.2 g methionine, 3.0 g phenylalanine, 1.2 g tryptophan, 1.8 g tyrosine, 1.2 g uracil, 9 g valine, 1.5 g aspartic acid, 1.5 g glutamic acid, 1.5 g threonine, 1.5 g serine, and 1.5 g proline) lacking histidine, arginine, and lysine + canavanine (50 mg/L) + thialysine (50 mg/L) and grown for 48 hours at 30°C (MATa cells are selected via *spHIS5* expression by the MATa-specific *MFA1* promoter while canavanine and thialysine select against unsporulated diploids). After 48 hours these plates were replica plated onto fresh plates containing the same media as before and grown for 24 hours at 30°Cin order to generate “tighter” spots for subsequent steps. Second, for selection of the knockout library gene deletion, the MATa strains were replica plated onto plates containing SD – His/Arg/Lys + canavanine + thialysine + G418 and grown for 48 hours at 30°C. Third, for selection of the *ded1-ΔCT* mutation, strains were replica plated onto plates containing SD – His/Arg/Lys + canavanine + thialysine + G418 + hygromycin and grown for 48 hours at 30°C. Growth on the final set of plates generated the DKO strains used for phenotypic screening.

### Phenotype Scoring

After generation of the double knockout strains, the strains were tested for growth fitness on media containing rapamycin. Both the single-mutant *ded1-ΔCT* query strain and some library deletion strains are resistant or sensitive to rapamycin (see Figure 1B, Supplemental Table S4, and (Aryanpur et al. 2019)). Therefore, DKO strains were scored by normalizing growth to that of the single-mutant parent strains via a multiplicative method (Baryshnikova et al. 2010). Fitness was assessed for each of the following on YPD and YPD + rapamycin (200 ng/mL) media: Y15583-13.2b (as a wild-type control), *ded1-ΔCT* query strain, single knockout library strains, and DKO strains. While still plated in a 384-well format, YPD plates were grown for 2 days and YPD + rapamycin plates were grown for 5 days at 30°C. Plates were scanned using a flat-bed scanner (Epson) and colony size was analyzed via SGA Tools (http://sgatools.ccbr.utoronto.ca), taking into account all four strains/controls in both conditions (Wagih et al. 2013). SGA Tools calculated fitness as the number of pixels contained in each spot for each strain. Then, using the single knockout in both conditions, the query strain in both conditions, and the wild-type control in both conditions as parameters for comparison, a normalized score was generated that represented the “interaction score” for the DKO strain on a scale from 0 to 2. This score thus provides a quantitative measure of the synthetic interaction between the *ded-ΔCT* allele and the library deletion for rapamycin-dependent growth, where 0 represents no growth of the double mutant, which is an extremely negative synthetic interaction (synthetic-lethal), 1 represents the expected growth given no interaction, and 2 represents much better growth on rapamycin than expected for the double mutant, a highly positive synthetic interaction (for technical reasons, the actual maximum was 1.961 rather than 2.00).

**Figure 1:**
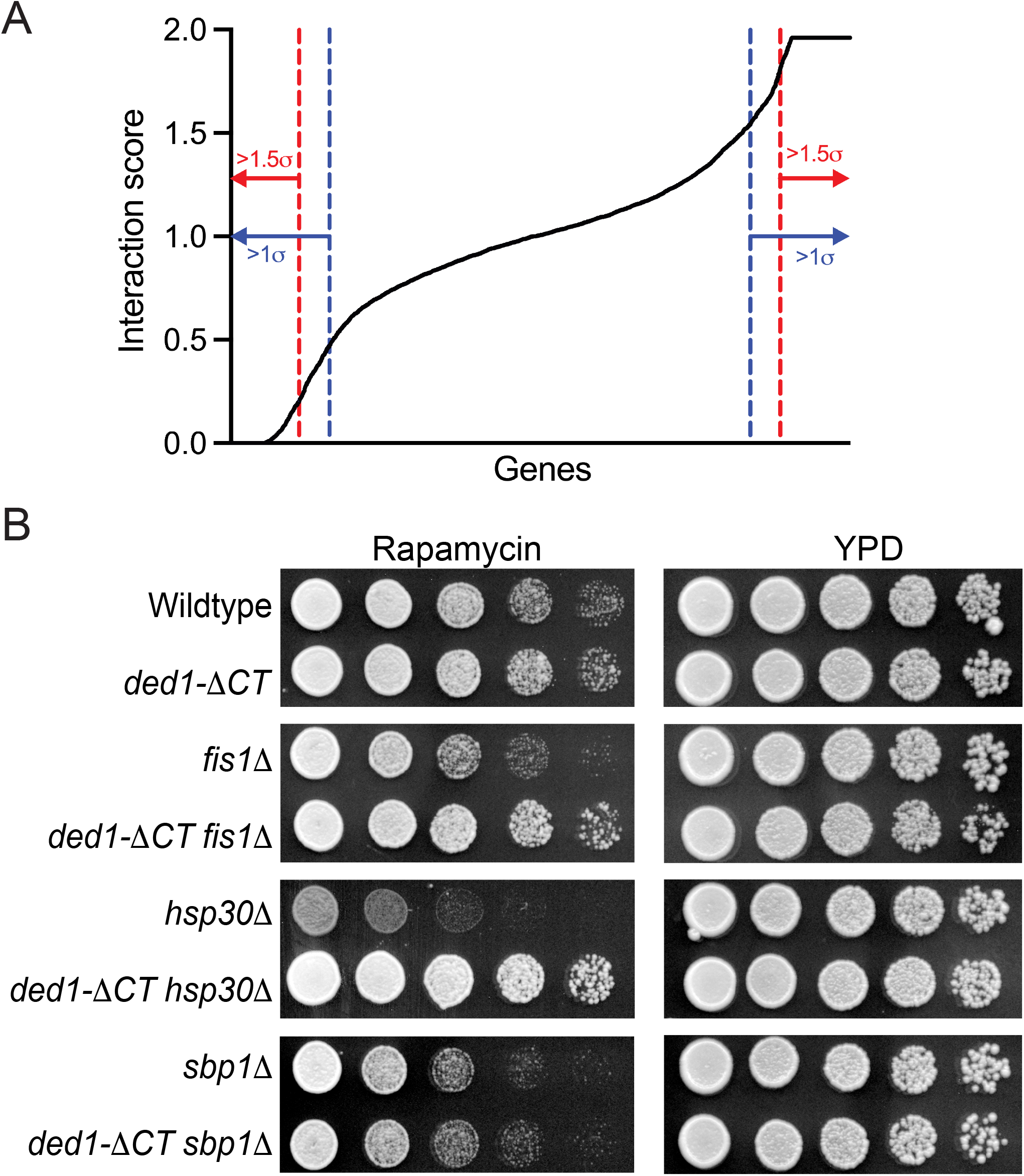
Identification of genetic interactions with *ded1-ΔCT* following rapamycin treatment. (A) The synthetic interaction scores with *ded1-ΔCT* for growth on rapamycin for all genes tested (4799) are shown in ascending order. Dashed blue lines represent 1 standard deviation from the mean score, and dashed red lines represent 1.5 standard deviations. Genes falling outside of these thresholds were considered “hits” and used for further analysis. (B) Growth of single (*fis1Δ, hsp30Δ*, and *sbp1Δ*) and *ded1-ΔCT* double mutants for three representative hits on rich media (YPD) and rich media plus rapamycin are shown. Five-fold serial dilutions were grown at 30°C for 2 (YPD) or 4 (Rapamycin) days. Synthetic interactions were observed consistent with the screen results.

### Classification and Verification of Hits

To determine cut-offs for further analysis of the hits, the interaction scores for all 4,799 DKO strains were analyzed via Graphpad Prism. The score distribution had a mean of 1.007 and a standard deviation of 0.534. Cut-offs were established at 1.0 and 1.5 standard deviations above and below the mean wherein interaction scores below the lower cut-off (0.473 or 0.205, respectively) were analyzed as negative / “sensitive” interaction hits and scores above the upper cut-off (1.542 or 1.809) were analyzed as positive / “resistant” hits. The 1.5 standard deviation cut-off, which yielded 529 negative and 544 positive hits, was used for most subsequent analysis; however, the 1.0 standard deviation cut-off, which yielded 763 negative and 780 positive hits, was used for generating lists of SGD-annotated phenotypes.

To experimentally verify the phenotypes, single and double mutants from a selected number of hits were isolated from the screen strains and individually tested via serial dilution growth assays on YPD and YPD + rapamycin plates at 30°C as previously described (Aryanpur et al. 2017). In addition, the list of hits includes a number of genes that were expected based on previous studies (e.g. translation factors).

### Network Maps and GO-Term Analysis

Network maps were generated with the hits (1.5 standard deviation cut-off) using STRING (https://string-db.org/) (Snel et al. 2000; Szklarczyk et al. 2021). STRING was then used to perform k-means clustering on the hits, where nodes (genes) were clustered into a predetermined number of clusters such that each node is related more strongly to other nodes within the same cluster than to nodes within other clusters. The gap statistic method was used to determine that five clusters should be used for each of the two datasets (sensitive and resistant hits) (Tibshirani et al. 2001). We then used STRING to associate GO-terms that are significantly enriched within each of the clusters. GO-terms were ranked for each cluster based on the “strength” feature in the STRING analysis, where strength is defined as log_10_ of the ratio between the number of proteins in the hits that are annotated with a specific GO-term and the number of proteins that would be expected to be annotated with this term in a random network of the same size. GO-term lists were manually curated by eliminating redundant or vague (“cytoplasm”) terms, and up to 10 GO-terms are shown. Complete lists of all significantly enriched GO-terms are included in the Supplemental Tables.

For the GO-term analysis of annotated hits (Table 3 and Supplemental Tables S4 and S5), the hits (1.0 standard deviation cut-off) were examined for a previously annotated phenotype of rapamycin (sirolimus) resistance or sensitivity using the “YeastMine” feature of the *Saccharomyces* Genome Database (SGD). Four categories of hits were thus generated: sensitive hits (as a double mutant with *ded1-Δ*CT) with a previously annotated rapamycin sensitivity (203 hits), sensitive hits with annotated rapamycin resistance (54), resistant hits with rapamycin sensitivity (78), and resistant hits with resistance (60). Note that only about one-quarter of the hits were annotated for a rapamycin phenotype in SGD; thus, three of the four categories were too small for effective network mapping. Instead, overrepresented GO-terms (biological process complete) were generated for each category using PANTHER (pantherdb.org) via Fisher’s exact test (Mi et al. 2021). GO-terms with less than 4 associated genes were deleted and then were curated as above with the 8 most-enriched terms shown in Table 3 (complete list in Supplemental Table S5).

## Results & Discussion

We crossed a *ded1-ΔCT* mutant to a knockout library of non-essential genes and assessed the resulting 4,799 double mutants for their growth on rapamycin-containing media. We then assigned each pair a synthetic interaction score (from 0 to 2, with 1 representing no interaction) after normalizing for growth on rapamycin of both single mutant parent strains, where a low score indicates that the double mutant grew less well on rapamycin than expected based on the single mutant phenotypes (a negative synthetic interaction), and a high score indicates that the double mutant grew better than expected (a positive synthetic interaction). The interaction scores were distributed across the range of possible scores with an overall mean of 1.007 (Figure 1A). A large number of mutants showed strong synthetic interactions with 266 scoring at the lowest value possible and 453 at the highest, respectively. Cut-offs for candidate hits were established as any mutants with an interaction score more than 1.0 or 1.5 standard deviations from the mean, depending on the downstream analysis (Figure 1A, Supplemental Table S1).

Results were verified by individually testing growth of selected hits. In Figure 1B, three examples are shown. The *fis1-null ded1-ΔCT* and *hsp30-null ded1-ΔCT* double mutants both grew significantly better on rapamycin than the *fis1-null* and *hsp30-null* mutants alone (positive interactions), while the *sbp1-null ded1-ΔCT* double mutant grew similarly to the *sbp1-null* mutant alone (and more poorly than the *ded1-ΔCT* single mutant), indicating a negative interaction. Thus, the growth assays agreed with the results from the screen, although we also found that some strongly negative interactions were also synthetic lethal or synthetic sick in the absence of rapamycin (data not shown). Further supporting the validity of the screen, genes expected to interact were also obtained, including yeast FKBP1 (*FPR1*) and numerous genes involved in translation (*GCN2*, ribosomal genes, etc.).

To organize the large number of hits from the screen, we conducted protein network analysis using STRING (Snel et al. 2000; Szklarczyk et al. 2021). An interaction network was built with all of the genes showing strongly negative synthetic interactions (“sensitive” hits) with *ded1-ΔCT* in the presence of rapamycin (529 genes with interaction score more than 1.5 standard deviations from the mean), and this network was then partitioned into 5 groups by *k*-means clustering (Figure 2A). A similar network was built and clustered for the 544 genes showing strongly positive synthetic interactions (“resistant” hits, Figure 2B). These clusters were then analyzed for Gene Ontology (GO) terms that are enriched in these subsets in order to determine which cellular processes and pathways interact with Ded1 most strongly during stress conditions.

**Figure 2:**
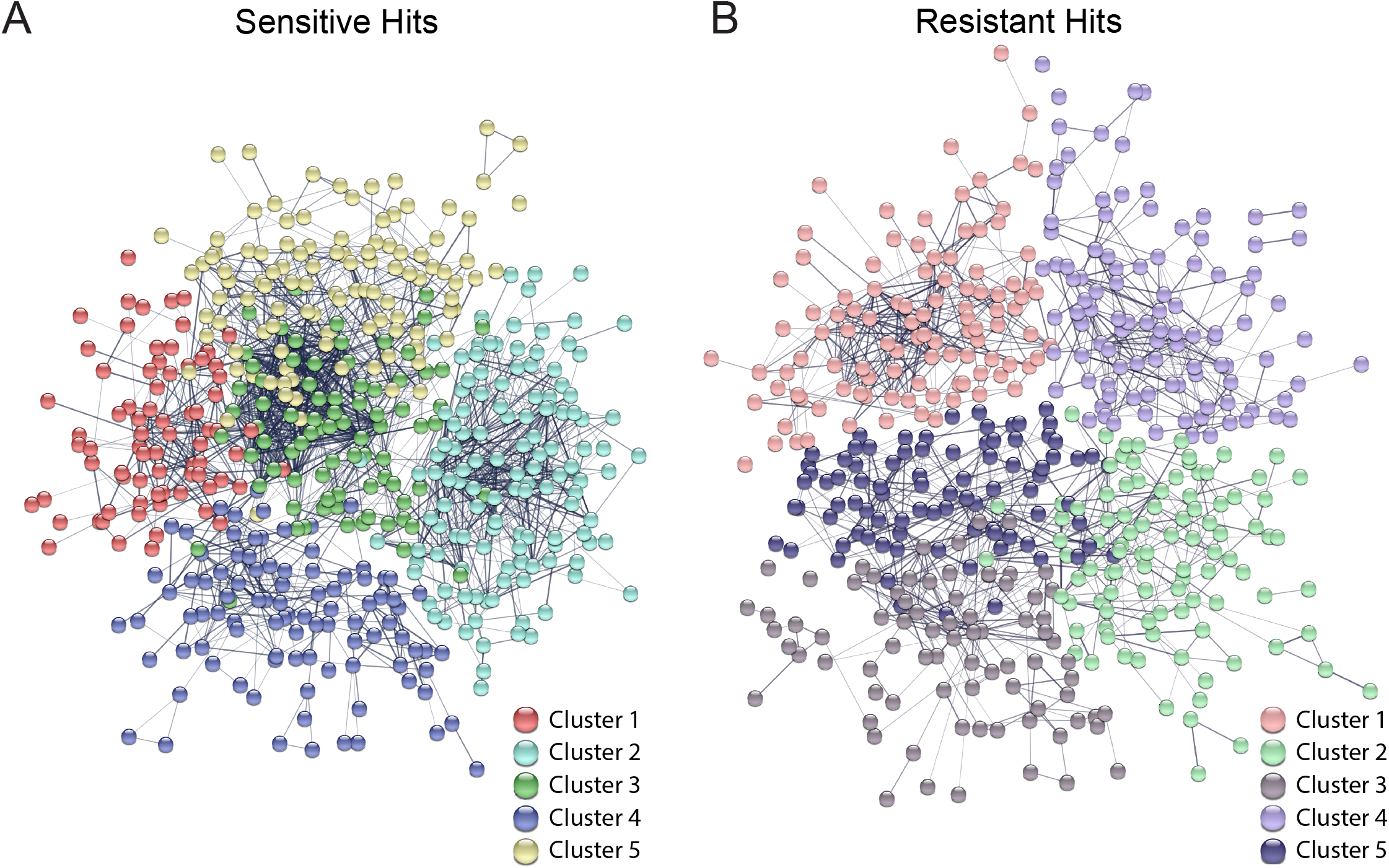
Network cluster maps of interacting genes. Network maps were generated using STRING for the synthetic negative / sensitive (A) and the synthetic positive / resistant (B) hits that exceeded the 1.5 standard deviation threshold below and above the mean interaction score, respectively. Thickness of the edges between genes signifies the strength of data support for the interaction. Disconnected nodes / genes are not shown. Clusters were generated via k-means clustering. The identity of each cluster (#1-5) is labeled below, and corresponding colors were used in Tables 1 and 2.

### Negative interactors / “sensitive” hits

Tables 1 and 2 show the most enriched GO-terms in each cluster (up to 10), following curation to remove highly similar terms (for complete lists of GO-terms, see Supplemental Tables S2 and S3). “Sensitive” cluster 1 included a substantial number of genes involved in amino acid metabolism, including multiple genes involved in synthesis of several different amino acids (arginine, isoleucine, serine, etc.) as well as synthesis of complex carboxylic acids (Table 1). Amino acid synthesis pathways are often upregulated in nutrient-poor conditions, so these genes may represent downstream targets regulated by Ded1, although amino acid availability also regulates TOR activity upstream (Crawford and Pavitt 2019; Saxton and Sabatini 2017). Cluster 1 also included eight genes involved in mitochondrial translation, which may affect energy production for translation. Sensitive cluster 2 yielded a number of GO terms that are related to membrane-mediated trafficking, including phosphatidylinositol signaling, intralumenal vesicle formation, endosomal transport, and vacuole regulation. These genes could be regulating Ded1 activity through its degradation along with its binding partner eIF4G1 during cell stress (Aryanpur et al. 2019; Kelly and Bedwell 2015), or they could be cross-talk from TOR-dependent regulation of autophagy and endosomal trafficking (Hatakeyama et al. 2019; Saxton and Sabatini 2017; Strahl and Thorner 2007). Cluster 2 also included eight genes involved in ATP export, which may again reflect an effect on energy production.

**Table 1:**
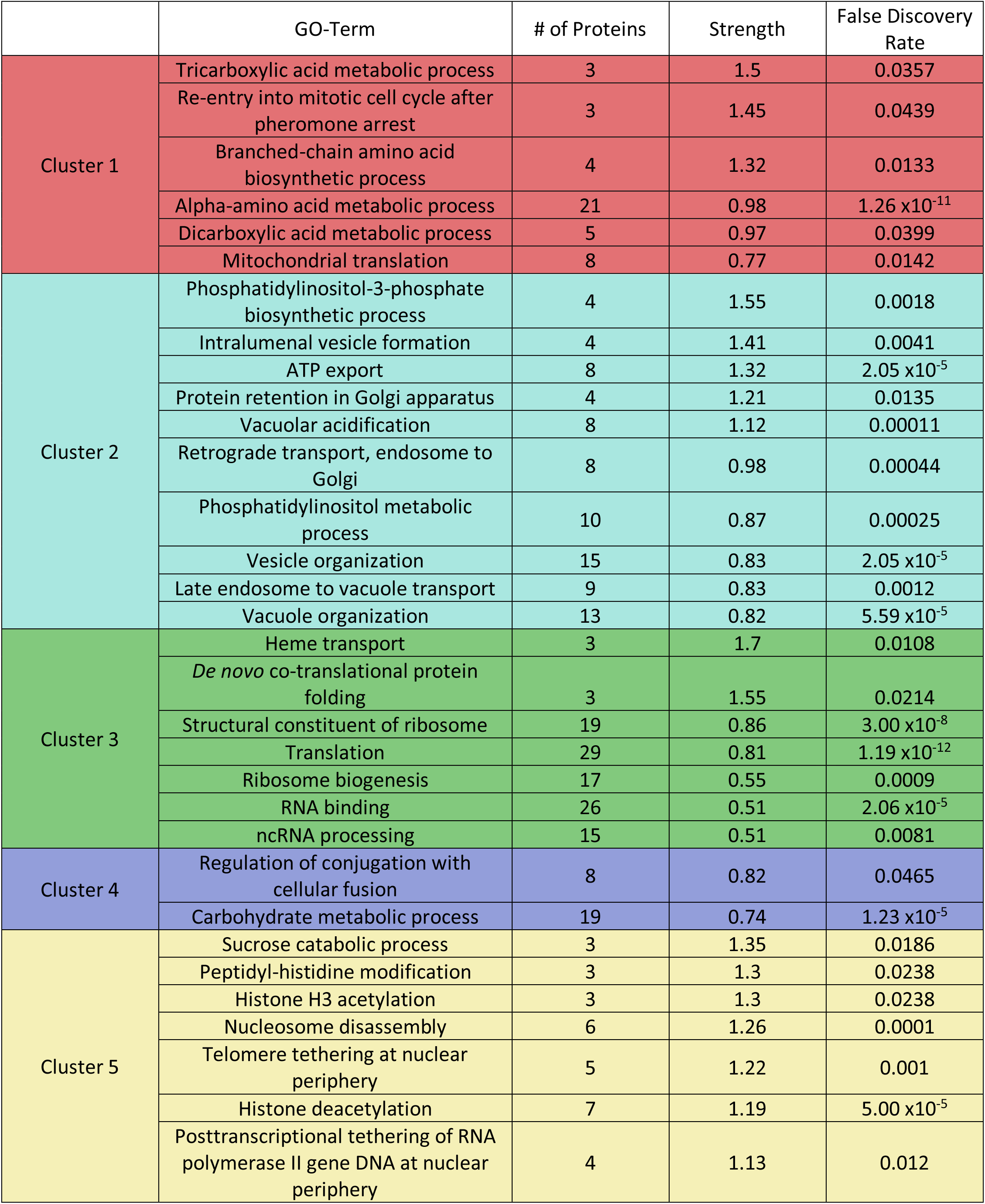
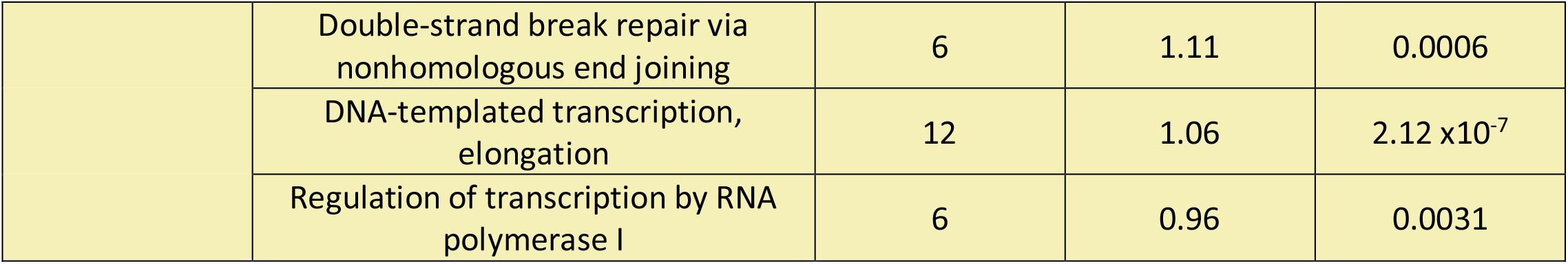
Enriched GO-terms in cluster analysis of sensitive hits

|  | GO-Term | # of Proteins | Strength | False Discovery Rate |
| --- | --- | --- | --- | --- |
| Cluster 1 | Tricarboxylic acid metabolic process | 3 | 1.5 | 0.0357 |
|  | Re-entry into mitotic cell cycle after pheromone arrest | 3 | 1.45 | 0.0439 |
|  | Branched-chain amino acid biosynthetic process | 4 | 1.32 | 0.0133 |
| | Alpha-amino acid metabolic process | 21 | 0.98 | $1.26 \times 10^{-11}$ |
|  | Dicarboxylic acid metabolic process | 5 | 0.97 | 0.0399 |
|  | Mitochondrial translation | 8 | 0.77 | 0.0142 |
| Cluster 2 | Phosphatidylinositol-3-phosphate biosynthetic process | 4 | 1.55 | 0.0018 |
|  | Intraluminal vesicle formation | 4 | 1.41 | 0.0041 |
| | ATP export | 8 | 1.32 | $2.05 \times 10^{-5}$ |
|  | Protein retention in Golgi apparatus | 4 | 1.21 | 0.0135 |
|  | Vacuolar acidification | 8 | 1.12 | 0.00011 |
|  | Retrograde transport, endosome to Golgi | 8 | 0.98 | 0.00044 |
|  | Phosphatidylinositol metabolic process | 10 | 0.87 | 0.00025 |
| | Vesicle organization | 15 | 0.83 | $2.05 \times 10^{-5}$ |
|  | Late endosome to vacuole transport | 9 | 0.83 | 0.0012 |
| | Vacuole organization | 13 | 0.82 | $5.59 \times 10^{-5}$ |
| Cluster 3 | Heme transport | 3 | 1.7 | 0.0108 |
|  | *De novo* co-translational protein folding | 3 | 1.55 | 0.0214 |
| | Structural constituent of ribosome | 19 | 0.86 | $3.00 \times 10^{-8}$ |
| | Translation | 29 | 0.81 | $1.19 \times 10^{-12}$ |
|  | Ribosome biogenesis | 17 | 0.55 | 0.0009 |
| | RNA binding | 26 | 0.51 | $2.06 \times 10^{-5}$ |
|  | ncRNA processing | 15 | 0.51 | 0.0081 |
| Cluster 4 | Regulation of conjugation with cellular fusion | 8 | 0.82 | 0.0465 |
| | Carbohydrate metabolic process | 19 | 0.74 | $1.23 \times 10^{-5}$ |
| Cluster 5 | Sucrose catabolic process | 3 | 1.35 | 0.0186 |
|  | Peptidyl-histidine modification | 3 | 1.3 | 0.0238 |
|  | Histone H3 acetylation | 3 | 1.3 | 0.0238 |
|  | Nucleosome disassembly | 6 | 1.26 | 0.0001 |
|  | Telomere tethering at nuclear periphery | 5 | 1.22 | 0.001 |
| | Histone deacetylation | 7 | 1.19 | $5.00 \times 10^{-5}$ |
|  | Posttranscriptional tethering of RNA polymerase II gene DNA at nuclear periphery | 4 | 1.13 | 0.012 |
|  | Double-strand break repair via nonhomologous end joining | 6 | 1.11 | 0.0006 |
| | DNA-templated transcription, elongation | 12 | 1.06 | $2.12 \times 10^{-7}$ |
|  | Regulation of transcription by RNA polymerase I | 6 | 0.96 | 0.0031 |

**Table 2:** Enriched GO-terms in cluster analysis of resistant hits

|  | GO-Term | # of Proteins | Strength | False Discovery Rate |
| --- | --- | --- | --- | --- |
| Cluster 1 | Ribosome | 16 | 0.51 | 0.0358 |
| Cluster 2 | Hydrolase activity, hydrolyzing O-glycosyl compounds | 8 | 1.01 | 0.0036 |
| Cluster 3 | Regulation of transcription from RNA polymerase II promoter in response to oxidative stress | 3 | 1.39 | 0.0449 |
|  | Peptidyl-tyrosine dephosphorylation | 4 | 1.17 | 0.0293 |
|  | Regulation of MAPK cascade | 5 | 1.04 | 0.0216 |
|  | Negative regulation of signal transduction | 6 | 1 | 0.0119 |
|  | Regulation of signal transduction | 12 | 0.83 | 0.00032 |
|  | Protein glycosylation | 7 | 0.83 | 0.0194 |
|  | Cellular response to abiotic stimulus | 6 | 0.8 | 0.0449 |
|  | Establishment or maintenance of cell polarity | 9 | 0.76 | 0.0096 |
| Cluster 4 | Aerobic respiration | 11 | 0.81 | 0.0083 |
| Cluster 5 | AP-type membrane coat adaptor complex | 5 | 1.39 | 0.0045 |
|  | Late endosome | 5 | 0.87 | 0.0499 |
|  | Vesicle | 12 | 0.53 | 0.0366 |
|  | Bounding membrane of organelle | 20 | 0.38 | 0.0366 |

Sensitive cluster 3 was largely focused on translation (Table 1), including 19 ribosomal proteins and at least 10 additional translation factors. Other aspects of translation were also represented, including ribosome biogenesis and non-coding RNA processing, which mostly consisted of tRNA processing genes (Supplemental Table S2). These are likely affecting the ability of Ded1 to regulate translation. Three protein chaperones that act co-translationally as well as three heme transport genes were also present in this cluster. Sensitive cluster 4 yielded relatively few GO terms, including the regulation of conjugation (mating) as well as carbohydrate metabolism. We have observed that *ded1-ΔCT* mutants have somewhat delayed sporulation compared to wild-type cells (data not shown), so these interactions are consistent with a Ded1 function in yeast mating/sporulation. Sensitive cluster 5 included a more diverse set of terms, including sucrose catabolism, peptidyl-histidine modification, double-strand break repair, and several terms related to chromatin remodeling and transcription. Alterations in chromatin state and/or transcription are of course part of stress responses and could represent upstream regulation or downstream targets of Ded1 activity following TOR inactivation (Crawford and Pavitt 2019; Saxton and Sabatini 2017). The three peptidyl-histidine modification genes all target translation factors for modification (ribosomal proteins and elongation factors), which may explain their genetic interaction with Ded1 (Al-Hadid et al. 2016; Uthman et al. 2013).

### Positive interactors / “resistant” hits

Despite a similar number of hits, the resistant clusters yielded fewer GO terms than the sensitive hits, suggesting a more diverse set of genes overall (Table 2 and Supplemental Table S3). Resistant cluster 1 included a number of ribosomal and ribosomal-related proteins, showing that alterations in different proteins involved in translation have the potential to either synergize or antagonize Ded1 function during cell stress. Resistant cluster 2 only yielded one significant GO-term, for hydrolase activity. These hydrolases are all involved in cell wall regulation during both sporulation and cytokinesis following mitosis. Their link to Ded1 is unclear but may be through sporulation and/or changes to the cell cycle during stress.

Cluster 3 gave the largest number of GO-terms for the resistant hits with particular enrichment for signal transduction genes, particularly the MAPK pathway. Notably, several of these (e.g. *PTC2, SDP1, PTP2*) are phosphatases that negatively regulate the MAPK pathway (Martin et al. 2005); therefore, their deletion would tend to increase growth and might synergize with increased growth in the *ded1-ΔCT* mutant in rapamycin. Other GO-terms in this cluster included transcriptional responses to oxidative stress, which fits well with Ded1 function in stress, and protein glycosylation, which has unclear links to Ded1. Only one GO-term, aerobic respiration, was associated with resistant cluster 4. This may again be due to energy requirements during stress. Finally, resistant cluster 5 included several GO-terms associated with membrane trafficking, similar to sensitive cluster 2, although with a more specific focus on vesicle trafficking, specifically. The relationship to Ded1 function is unclear, although these interactions may be due to TOR-dependent changes in membrane trafficking that affect Ded1 activity during stress.

### Annotation of hits by rapamycin-dependent phenotype

In theory, “resistant” synthetic interactions in this screen could be generated either by synergistic effects of a rapamycin-resistant mutation and the rapamycin-resistant *ded1-ΔCT* allele (a “resistant-resistant” phenotype), or by suppression of a rapamycin-sensitive mutation by *ded1-ΔCT* (a “resistant-sensitive” phenotype). Likewise, “sensitive” interactions could be due to suppression of *ded1-ΔCT* rapamycin resistance by a rapamycin-sensitive mutation (“sensitive-sensitive”), or by suppression by a rapamycin-resistant mutation (“sensitive-resistant”). To attempt to assign hits to these various categories, we mined the phenotypes of the *Saccharomyces* Genome Database for those genes with mutations annotated as resistant or sensitive to rapamycin, and then we correlated these with the genes in our screen with an interaction score more than 1.0 standard deviation from the mean. Only a minority of the hits were annotated for a rapamycin-dependent phenotype (138 resistant and 257 sensitive hits), so we were not able to conduct in-depth network analyses for these subsets. Nonetheless, we generated enriched GO-terms for each of the four subsets, which are summarized in Table 3 (for complete lists, see Supplemental Tables S4 and S5). The “resistant-resistant” subset included GO-terms for regulation of GTPase activity, regulation of transcription, ribosomal biogenesis, and response to stress. The resistant-sensitive subset included transcription elongation, phosphatase activity, membrane trafficking, translation, and DNA repair. The sensitive-sensitive subset included the largest number of annotated hits (203) and yielded the largest number of GO-terms, including intralumenal vesicle formation, other membrane trafficking terms, ATP export, TOR signaling, and DNA maintenance. By contrast, the sensitive-resistant category was the smallest with 54 genes, and GO-terms included transcription regulation, ubiquitination, and ion homeostasis. Overall, the GO-terms in this analysis largely corresponded to the terms from the cluster analysis above, with several new terms such as GTPase activity and ubiquitination. However, this division into subcategories may be useful in designing follow-up experiments to directly examine these interactions with Ded1.

**Table 3:** Enriched GO-terms in previously-annotated hits

|  | GO-Term | # of Proteins | Strength | P-value |
| --- | --- | --- | --- | --- |
| Resistant hits with known rapamycin resistance | Positive regulation of GTPase activity | 4 | 1.08 | 0.0005 |
|  | Negative regulation of transcription by RNA polymerase II | 6 | 0.64 | 0.0026 |
|  | Response to oxidative stress | 5 | 0.63 | 0.0066 |
|  | Regulation of intracellular signal transduction | 4 | 0.62 | 0.0167 |
|  | Ribosomal large subunit biogenesis | 4 | 0.51 | 0.0360 |
|  | Telomere organization | 4 | 0.51 | 0.0360 |
|  | Chromatin organization | 9 | 0.50 | 0.0022 |
|  | Positive regulation of transcription by RNA polymerase II | 9 | 0.50 | 0.0022 |
| Resistant hits with known rapamycin sensitivity | Positive regulation of DNA-templated transcription elongation | 4 | 0.85 | 0.0029 |
|  | Dephosphorylation | 4 | 0.77 | 0.0054 |
|  | Endocytosis | 6 | 0.67 | 0.0021 |
|  | Protein localization to membrane | 5 | 0.52 | 0.0188 |
|  | Regulation of translation | 6 | 0.49 | 0.0137 |
|  | Autophagy | 5 | 0.42 | 0.0419 |
|  | DNA repair | 8 | 0.41 | 0.0136 |
|  | Vesicle-mediated transport | 11 | 0.38 | 0.0059 |
| Sensitive hits with known rapamycin resistance | Negative regulation of transcription by RNA polymerase II | 9 | 0.90 | $2.03 \times 10^{-6}$ |
|  | RNA catabolic process | 5 | 0.64 | 0.0058 |
|  | Protein ubiquitination | 4 | 0.62 | 0.0168 |
|  | Positive regulation of transcription by RNA polymerase II | 8 | 0.53 | 0.0024 |
|  | Cellular ion homeostasis | 4 | 0.49 | 0.0423 |
| Sensitive hits with known rapamycin sensitivity | Intralumenal vesicle formation | 4 | 1.25 | 0.0003 |
| | ATP export | 9 | 1.22 | $4.63 \times 10^{-8}$ |
| | Protein localization to Golgi apparatus | 8 | 1.07 | $1.91 \times 10^{-6}$ |
|  | Maintenance of DNA trinucleotide repeats | 4 | 1.05 | 0.0010 |
|  | Positive regulation of TOR signaling | 4 | 0.98 | 0.0016 |
|  | Homoserine metabolic process | 4 | 0.92 | 0.0025 |
| | Retrograde transport, endosome to Golgi | 11 | 0.89 | $7.47 \times 10^{-7}$ |
| | Vacuolar acidification | 7 | 0.89 | $8.31 \times 10^{-5}$ |

In this screen, we obtained a large number of potential interactions with the *ded1-ΔCT* mutant. Many of these fell into categories that would be predicted by the known functions of Ded1 during cellular stress, including ribosomal proteins and translation factors, amino acid biosynthesis genes, and transcription and chromatin remodeling factors. Several more have more tangential links to the stress function of Ded1 and may represent cross-talk between cellular processes or even mRNAs that are translationally regulated by Ded1. These include genes involved in membrane trafficking, mitochondrial / energy production genes, and sporulation genes. Follow-up experiments to explore the links between Ded1 and these processes could lead to better understanding of the coordination and regulation of cellular stress responses. Lastly, some hits were in unexpected categories, such as cell wall hydrolases, protein glycosylation, and GTPase activity. Future work may be able to elucidate these interactions with Ded1 and how they contribute to stress or other cellular responses.

## Data Availability Statement

Strains and plasmids are available upon request. The supplemental tables contain complete lists of all data and analysis from the screen, including interaction scores for all genes (Supplemental Table S1), SGD-annotated phenotypes (S4), and complete lists of GO-terms (S2, S3, and S5).

## Acknowledgments

We would like to thank Matt Kaplan, director of the University of Arizona Functional Genomics Core, for equipment use, training, and reagents. We would also like to thank members of the Bolger laboratory, past and present, for discussions and suggestions on the screen results and the manuscript.

